# Who are my Neighbours? Selection Targets Robustness to Full Replication Events, Not to Individual Mutations

**DOI:** 10.64898/2026.09.24.751893

**Authors:** Juliette Luiselli, Guillaume Beslon, Pieter van den Berg

**Affiliations:** Evolutionary Modelling Group, Department of Biology, KU Leuven, Leuven, 3000, Belgium; Centre of Microbial and Plant Genetics, KU Leuven, Leuven, 3000, Belgium; BioTiC Team, Inria Lyon La Doua, Villeurbanne, 69100, France; INSA-Lyon, INRIA, CITI EA3720, Lyon F-69621, France

**Keywords:** Fitness Landscape, Genome Size, Mutational Robustness, Replicative Robustness, Evolution

## Abstract

Theory predicts that populations should evolve mutational robustness — the ability to maintain a constant phenotype despite mutations. However, selection does not operate on individual mutations but on phenotypes resulting from entire replication events, which may involve a variable number of mutations. We therefore distinguish mutational robustness from replicative robustness, defined as the probability that a replication event produces similarly fit offspring. Mutational and replicative robustness may seem closely aligned at first sight, but we show with an evolutionary simulation model that they can be selected in opposite directions. Our results show that while genomes with high redundancy and/or large amounts of non-coding DNA are mutationally robust, they are replicatively fragile because their size increases the number of mutations per replication. This is especially relevant when genomes are at risk of severe disruption from chromosomal rearrangements. Our results expose limits of the fitness landscape metaphor in the context of robustness evolution: neutral plateaus may be mutationally robust but replicatively unstable and therefore disfavoured by selection, whereas narrow peaks arising from densely encoded genomes can replicate more reliably. Our results identify replicative robustness as a distinct target of selection that depends not only on tolerance to individual mutations but also on their overall likelihood of occurring.

## Introduction

Fitness landscapes are abstract representations of the relationship between biological variation and reproductive success, often visualised as a topographical surface to help understand the evolutionary trajectories of populations over time. Visualisation of fitness landscapes was first introduced by Sewall Wright in 1932 (Wright, 1932), and has since encouraged significant development in the field of evolutionary biology (De Visser and Krug, 2014; Fragata et al., 2019), both through the intuition it provides on evolving populations and through the theoretical insights obtained by the formal mathematisation of the concept (Pitzer and Affenzeller, 2012). Fitness landscapes can be defined at different levels, including genotype-to-fitness, phenotype-to-fitness, and molecular landscapes (Schuster and Stadler, 2023); here, we focus on genotype-to-fitness maps in stable environments.

Theory predicts that, under some conditions, selection can lead populations to occupy flatter regions of the fitness landscape rather than sharp peaks (‘survival of the flattest’ (Wilke et al., 2001; Wilke and Adami, 2003; Codoñer et al., 2006)). On such plateaus, populations can accumulate mutations without substantial loss of fitness, a property known as mutational robustness. This is especially relevant in asexual populations evolving under high mutation rates. At low mutation rates, selection is dominated by differences in immediate fitness, and any advantage associated with robustness is typically too weak to overcome these differences. As mutation rates increase, however, the fitness consequences of mutations become more prominent, and genotypes whose mutational neighbourhood contains a higher fraction of neutral or mildly deleterious variants can achieve a higher long-term reproductive success (Wilke et al., 2001; Lauring et al., 2013). As asexual species lack recombination, an individual’s offspring continue to explore the same region of the fitness landscape rather than being shuffled onto new genetic backgrounds, so that a lineage’s mutational neighbourhood - and hence its level of robustness - is transmitted intact, and the benefits accrue within the lineage. As a consequence, it has been argued that selection for mutational robustness can lead to genetic architectures that give rise to flatter fitness peaks (Wilke and Adami, 2003), such as high-redundancy architectures which can arise through gene duplication (Wagner, 2008a; Fares, 2015).

This perspective, however, relies on the premise that single mutations constitute the relevant steps on the fitness landscape and that mutational distance adequately captures proximity between genotypes. While the metaphor of the fitness landscape has been heavily debated (Kaplan, 2008; McCandlish, 2011; Zagorski et al., 2016), both its proponents and its critics typically retain mutational distance as the fundamental measure of proximity between genomes: evolutionary change is conceptualised as movement across the landscape through single-mutation steps. However, as most mutations occur during reproduction, selection acts on the outcome of *replication events*, which may entail a variable number of mutations, rather than on individual mutations themselves. While considering mutations one by one can be an appropriate proxy when considering only point mutations at low rates, this approximation may become problematic as the mutation rate increases, since multiple mutations are then more likely to occur during the same replication event, or when a mutation changes the genome size, as this can impact the probability of future mutations. Additionally, the number of mutations bundled into a replication cannot be treated independently of their fitness effects: genomes with more mutations per replication event may have milder mutations. For instance, increases in the amount of non-coding DNA decrease the average impact of individual mutations, but simultaneously increase the per-genome mutation rate, as there are now more base pairs that can mutate (Luiselli et al., 2025). From a quantitative genetics perspective, this means that mutational variance (a compound of both mutation rate and mutational effect size) can be impacted by mutations that alter genome size and redundancy (such as deletions and duplications) in intricate ways. While previous work has independently investigated the evolution of mutation rates and the evolution of mutational effects (*i*.*e*., mutational robustness), they have rarely been considered in concert (Lynch et al., 2016; LaBar and Adami, 2017).

Here, we present the results of a computational model of genome evolution showing that selection acts on *replicative robustness* (defined as the probability that an offspring has the same fitness as its parent after a full replication of the genome) rather than on *mutational robustness* (defined as the probability that a genome retains its fitness following a random mutation). Our results also show that this difference matters for genome size evolution. Specifically, our model shows that selection for replicative robustness favours genome reduction (thereby reducing the number of mutations generated during replication), even though this reduces mutational robustness because mutations are more likely to disrupt essential functions in short genomes which typically lack redundancy. These results are consistent with a different conceptualisation of the fitness landscape metaphor, where the steps between neighbouring genotypes reflect replication events rather than single mutations. As our results indicate, at least some plateaus in the mutational fitness landscape correspond to peaks in the replicative fitness landscape. Conversely, some peaks that appear unstable in the mutational fitness landscape are highly stable in the replicative fitness landscape, behaving more like plateaus.

## Materials and methods

To carry out this study, we developed an individual-based C++ simulator of genome evolution. A population of individuals, each carrying a genome, competes at each generation to replicate, and each replication can entail zero, one, or several mutations.

### Genome model

For each experiment, we follow a population of fixed size *N*. As depicted in Figure 1a, each individual possesses a single circular genome, on which there are genes, separated by non-coding regions of varying size. We consider a set of *g* different genes, each essential for replication. Each gene has the same size *l*. An individual must own at least one copy of each gene of the set, while additional copies do not affect its fitness. The fitness *w* of individual *i* is thus defined as

**Figure 1.**
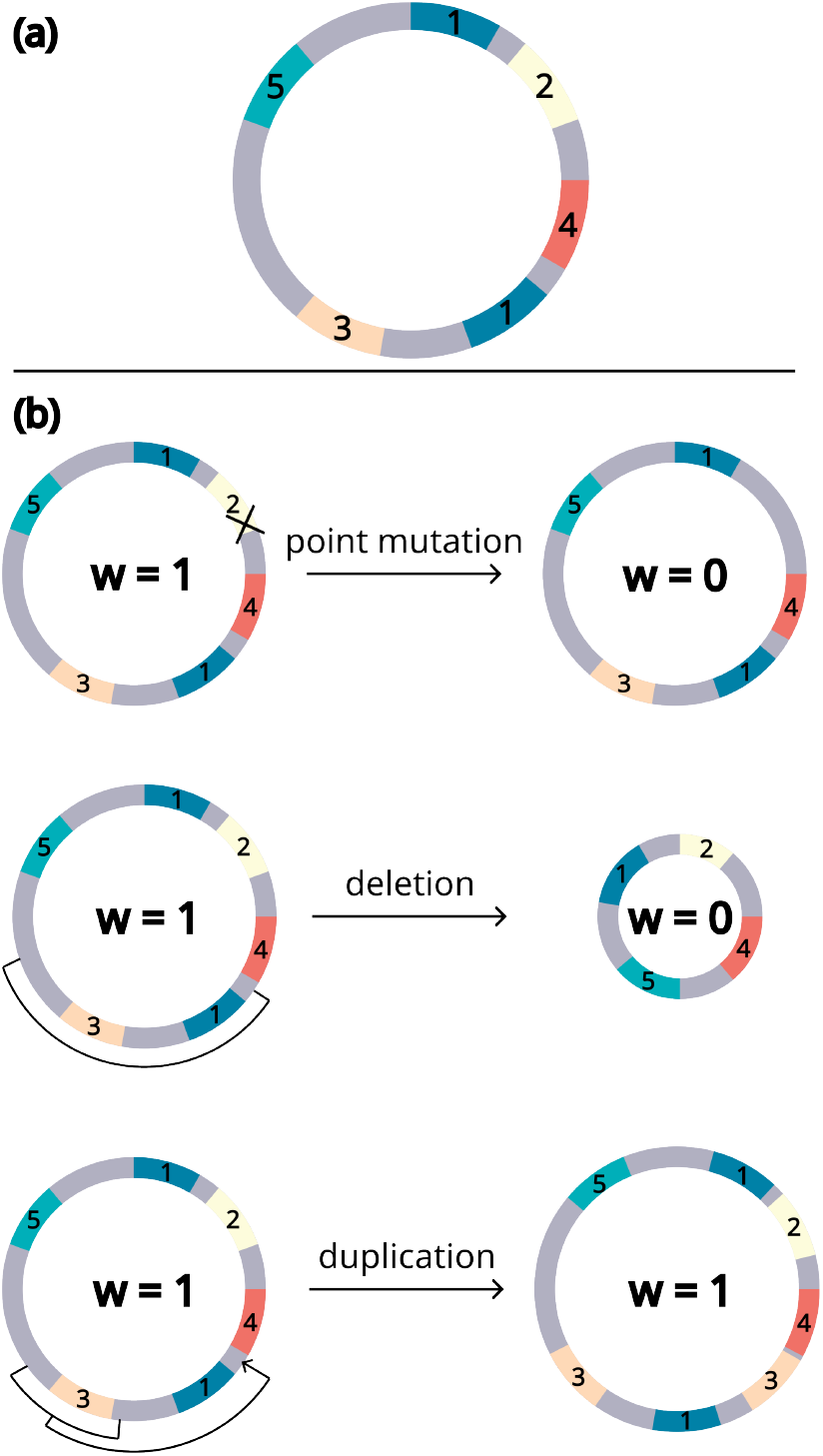
(a) Schematic representation of a genome in the model. Genes are all of the same size and are potentially separated by non-coding sections of varying sizes. The non-coding sections are in light grey, while the genes are in different colours based on their IDs. This genome has two copies of gene 1, and a single copy of genes 2, 3, 4 and 5. (b) Schematic representation of the possible mutation types, and their consequences on gene count and fitness. A point mutation within a gene renders it non-functional, turning it into a non-coding part of the genome. A deletion can delete one or several genes and is frequently lethal. A duplication can make new copies of genes.

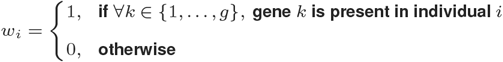

At each generation, all individuals are replaced by drawing (with replacement) *N* viable individuals (with a fitness of 1) as reproducers to populate the next generation. Consequently, some individuals leave no offspring, whereas others leave multiple offspring. Each reproducer is then replicated and possibly undergoes mutations, allowing genome structures to evolve over generations. Non-viable individuals do not reproduce.

### Mutations

During each replication event, mutations can occur. There are three mutation types, depicted in ??b: point mutations, duplications, and deletions. If a mutation disrupts a gene in any way (point mutation in a gene, duplication inserting within a gene, deletion of any part of a gene), then that gene is rendered non-functional, and its remnants become non-coding. Note that genes can be duplicated, which increases mutational robustness as it decreases the risk of future mutations being deleterious (disrupting one of the two gene copies will not affect fitness). Also note that the non-coding genome can vary freely: it can be duplicated or deleted without any impact on fitness. To reduce computational load, an upper bound of 10^9^ *bp* per individual is enforced, but this limit is rarely reached and does not impact our results (see Supp. Mat section S5).

Each type of mutation has a constant per-base mutation rate, and mutation breakpoints are uniformly distributed along the genome. The size of structural mutations (duplications or deletions) follows a geometric distribution of parameter 10*/L* where *L* is the total genome size. This means that larger genomes are more susceptible to larger mutations, which is a reasonable assumption as double-strand breaks forming rearrangements are more likely to be further apart in a larger genome. This is supported by the fact that chromosomal rearrangements have been documented to be larger in species with larger genomes (Cui et al., 2012; da Silva et al., 2019; Wellenreuther and Bernatchez, 2018).

### Simulation Design

Using this model, we evolve populations of different sizes (1, 000, 2, 000, 5, 000, 10, 000, and 20, 000 individuals), with different point mutation rates (5 × 10^−7^, 1 × 10^−6^, 1.5 × 10^−6^, 2 × 10^−6^, 5 × 10^−6^) and different deletion rates (2 × 10^−7^, 3 × 10^−7^, 4 × 10^−7^, 5 × 10^−7^ per base). The duplication rate is fixed at 1 × 10^−7^ per base. For each tested parameter combination, we run 50 replicates with independent random seeds. Each population is initially filled with clonal individuals that have one copy of each of the *g* = 50 different genes (of size *l* = 1, 000 bp each). Note that all parameter combinations include a mutational bias towards deletions, which are more likely than duplications — as is generally the case in the microbial world (Kuo and Ochman, 2009).

To evaluate the outcome of our experiments, we compare the average number of gene copies per individual in the common ancestors of the final populations, as well as the total genome size and the sizes of the coding and non-coding parts of the genome. We run simulations for 20, 000 generations. To reduce noise, which is typically caused by transient large neutral duplications, each data point is computed by averaging over the ancestors from generations 14, 000 to 15, 000. This is both late enough in the simulations to ensure equilibrium (see the temporal data in Supplementary Materials section S1: Figure S1, Figure S2 and Figure S3), and early enough compared to the 20, 000 generation to ensure coalescence. Taking the ancestor of the population rather ensures that we look at genomes that eventually get fixed in the population and allows for a straightforward comparison of populations of different sizes.

We also quantify the local flatness of the fitness landscape, *i*.*e*. the proportion of neighbours that have the same fitness, using the two definitions used in this study. Mutational robustness is defined as the probability that a genome retains full fitness following a random mutation sampled according to the underlying mutation spectrum (*i*.*e*., weighted by the relative mutation rates of the different mutation types). For each individual of interest, this probability is estimated using 100, 000 single mutation events of each type. Replicative robustness is defined as the probability that a genome retains full fitness after a replication event and is estimated by producing 100, 000 offspring per individual of interest. In this case, we use the same per-base mutation rates as in the corresponding evolutionary experiments.

### Statistical analysis

We used ordinary least squares (OLS) regression to model gene copy number as a function of mutation rates (Figure 2) or of population size (Figure 3), using Python statsmodels package (version 0.14.1). We used generalised linear models with Beta error structure (Ferrari and Cribari-Neto, 2004) to model the continuous response of mutational robustness and replicative robustness (both bound within [0, 1]) to changes in mutation rates (Figure 2) or population sizes (Figure 3), using Python `statsmodels` package (version 0.14.1). Error bands in all graphs show 95% confidence intervals for the expected mean. For models that included population size as a predictor, we used the logarithm of population size to account for the wide range of tested values.

**Figure 2.**
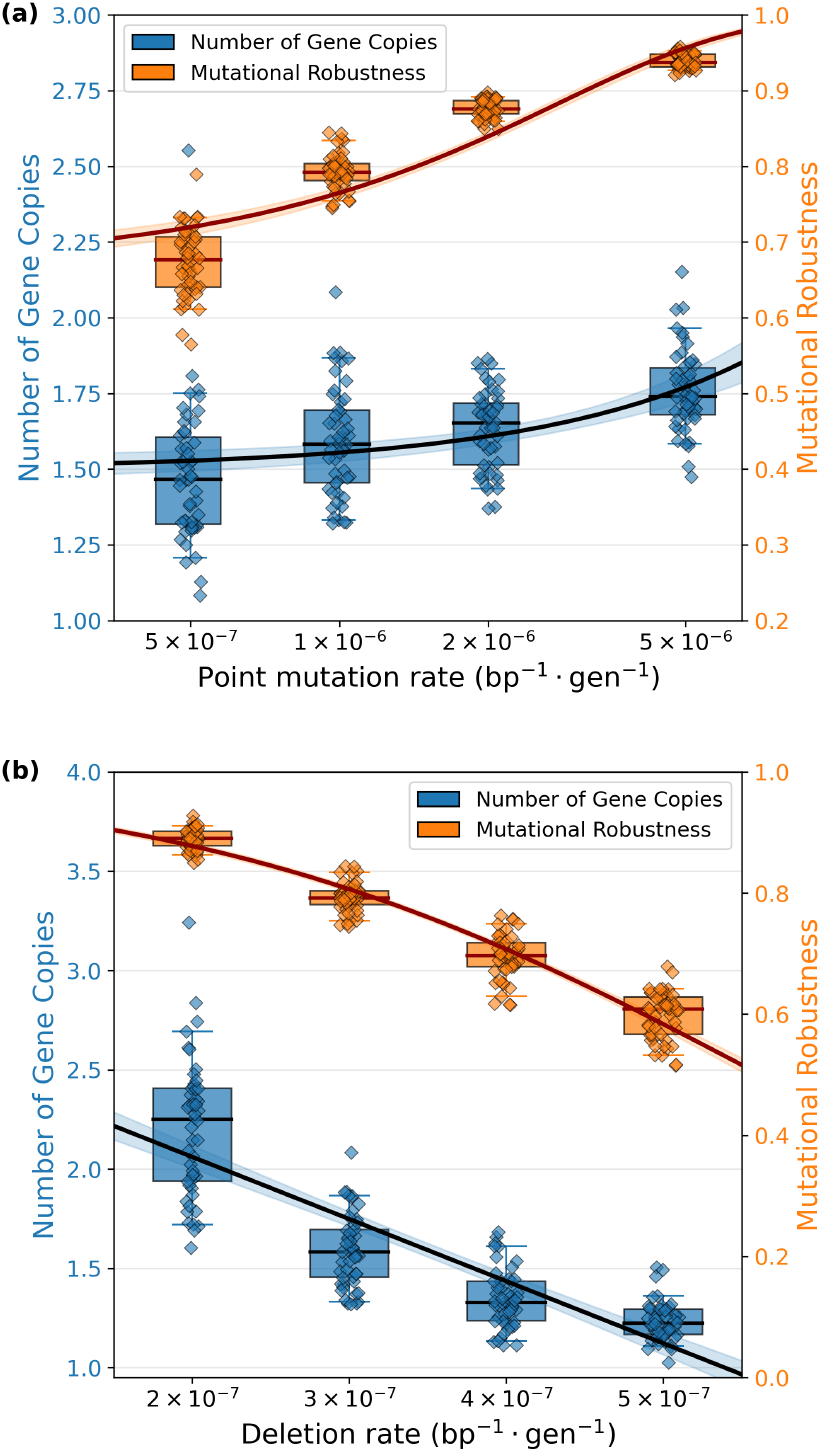
(a) Average number of gene copies (blue) and mutational robustness (orange) at equilibrium, for experiments with different point mutation rates. The other parameters (duplication rate = 1 × 10^−7^, deletion rate = 3 × 10^−7^, as well as population size *N* = 10, 000) are kept constant. A slight jitter has been added to the points to increase readability. Note that the scale of point mutation rate here is logarithmic to improve readability. (b) Average gene copy number (blue) and mutational robustness (orange) at equilibrium for experiments with different deletion rates. The other parameters (duplication rate = 1 × 10^−7^, point mutation rate = 1 × 10^−6^ and population size *N* = 10, 000) are kept constant. A slight jitter has been added to the points to increase readability.

**Figure 3.**
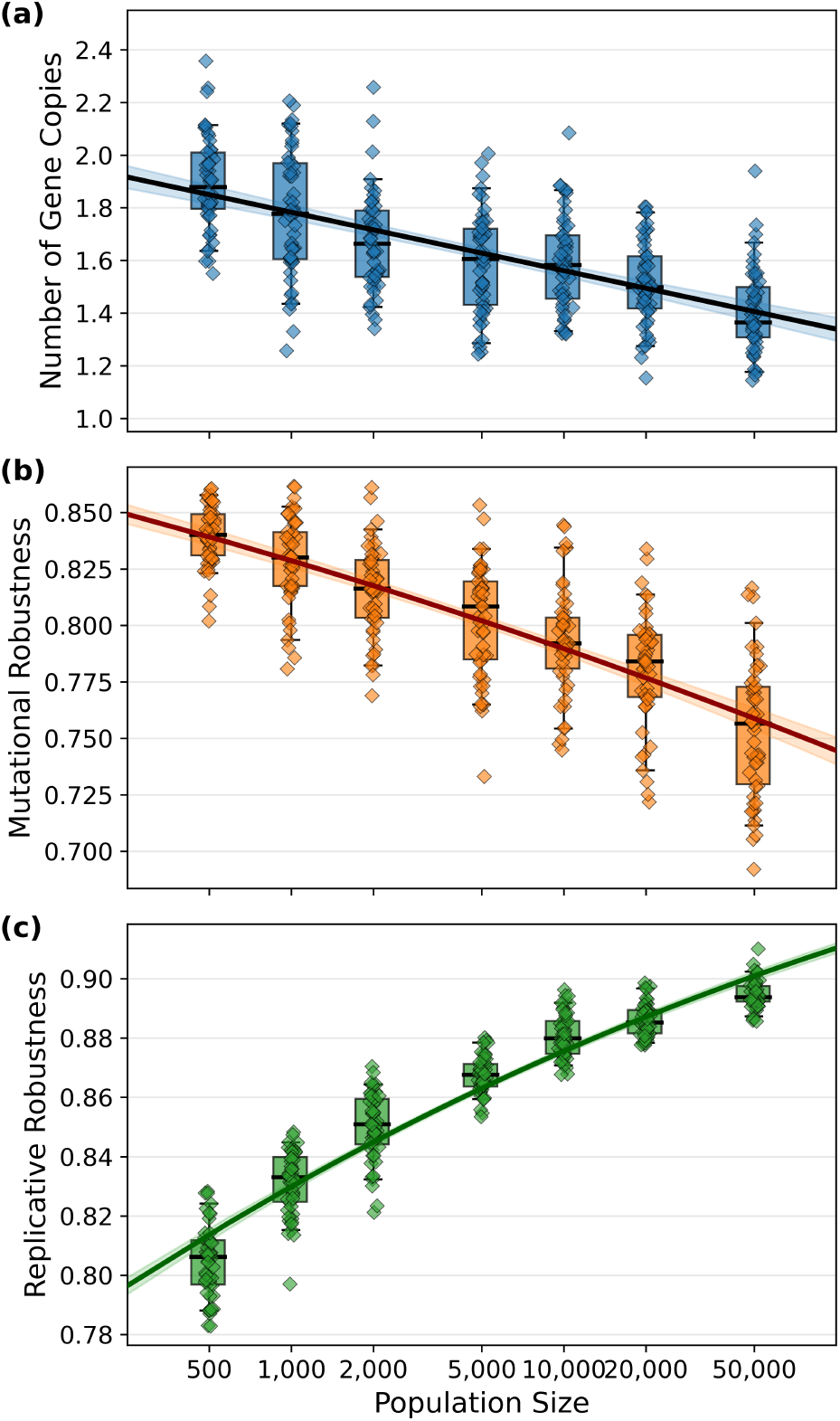
Average (a) number of gene copies (b) mutational robustness and (c) replicative robustness at equilibrium, for various population sizes. The other parameters are fixed at: duplication rate = 1 *×* 10^−7^, deletion rate = 3 *×* 10^−7^, point mutation rate = 1 *×* 10^−6^.

## Results

### Higher point mutation rates select for increased mutational robustness

The classical theory of survival of the flattest (*i*.*e*., selection of mutational robustness) predicts that higher rates of point mutations select for increased mutational robustness (Hulst et al., 2025; Bornholdt and Sneppen, 2000; Elena et al., 2007). In our model, this should result in more gene copies: while having several copies of a gene does not impact fitness directly, it does reduce the risk of losing the gene due to a mutation. Hence, in our model, having more gene copies is equivalent to being on a flatter region of the fitness landscape (*i*.*e*., a larger plateau).

Figure 2a shows that (at equilibrium) populations evolve more gene copies (linear regression, *p <* 0.001) and higher mutational robustness (beta GLM, *p <* 0.001) as the point mutation rate increases, with all other parameters held constant. The temporal data for these simulations—and for all other results presented in the manuscript—are available in the Supplementary Materials section S1. Note that the increase in mutational robustness is not only due to the increase in gene copy number, but also to the increase in non-coding genome size due to the high rate of pseudogenization induced by point mutations (see details in Figure S4). This aligns with the predictions from the theory that evolution tends to bring populations towards large fitness plateaus, especially when the mutational pressure is high. However, as we will discuss below, this pattern is quite specific for point mutations: structural mutations can have quite different effects on genome evolution.

### Higher deletion rates select against mutational robustness

One might naively expect that deletions, which have an even higher disruptive potential than point mutations, would also select for high redundancy (*i*.*e*., high number of gene copies) to increase robustness and reduce their disruptive impact. However, our simulations show that this is not the case. On the contrary, Figure 2b shows that (at equilibrium), populations evolve fewer gene copies and lower mutational robustness as the deletion rate increases, with all other parameters held constant. The reason for this could be that even though increasing gene copy number protects genes against mutations through redundancy, it also puts nearby (potentially non-duplicated) genes at a higher risk of being distorted by deletions. As the deletion rate increases, populations thus evolve toward increasingly narrow peaks (*i*.*e*. less flat regions) of the fitness landscape. Hence, different mutational loads (here, from point mutations or from deletions) result in different genome architectures.

The reduction in the number of gene copies under an elevated deletion rate is expected to be partly driven by the mutational bias towards deletions, but selection might either oppose or reinforce this tendency. To disentangle the contribution of selection from that of a mutational bias, we next compare the gene copy number evolved at equilibrium across populations of different sizes under relatively high deletion rates. Because selection is more effective in larger populations, we expect stronger effects of selection on gene copy number in larger populations. If selection favours genome reduction, larger populations should evolve fewer gene copies; and conversely, if selection favours the maintenance of additional copies, larger populations should evolve more gene copies. In contrast, if mutational bias were the only force shaping genome size, gene copy number should be independent of population size. Figure 3A shows that the former is the case: larger populations evolve lower gene copy numbers, indicating that genome reduction is driven by selection.

Importantly, this reduction has opposing effects on mutational and replicative robustness. While this reduction in gene copy number decreases mutational robustness (any single mutation is *more* likely to reduce fitness; Figure 3B), it increases replicative robustness (full replications are *less* likely to reduce fitness Figure 3C). This exposes a trade-off between mutational and replicative robustness under these circumstances: by evolving smaller genomes, populations reduce the number of mutations per generation, but increase the average fitness impact of any individual mutation. This pattern holds across a wide range of point mutation and deletion rates (see Figure S8 and Figure S9). Indeed, if we examine the mutational and replicative robustness of engineered genomes that have constant fractions of coding DNA but varying size, we find the same patterns: mutational and replicative robustness both increase with genome size when only point mutations are accounted for, while for deletions mutational robustness increases with genome size but replicative robustness decreases with genome size (see Figure S11).

The evolutionary dynamics described above depend on how the overall mutation rates scale with genome size. In our model, mutation rates are defined per base, such that the per-genome mutation rate increases with genome size. Many previous theoretical studies of robustness evolution have reported that robustness selection leads to an increase in gene copy number, or genome size more generally (Osterwalder et al., 2018; Posadas-García and Espinosa-Soto, 2022; Ebadi et al., 2023). However, few of these studies allow for structural mutations or genome size variation. When duplications and deletions are included, they typically concern single genes rather than genomic segments, and mutation rates are often assumed to be fixed per genome (Bornholdt and Sneppen, 2000; Elena et al., 2007; LaBar and Adami, 2017). To assess the impact of this assumption, we implemented a variant of our model in which mutation rates are fixed per genome rather than per base. Note that the rest of the model is unchanged. In particular the average mutation size still depends on the genome size.

#### This modification of our model radically alters the evolutionary outcome

fixed per-genome mutation rates lead to a rapid increase in gene copy number (quickly reaching 100 copies; Figure 4A). This extremely high copy number confers near-perfect robustness (both mutational and replicative, seeFigure 4B and C). We also observe a slight increase in the number of copies with increasing population sizes (see Supp Mat. Figure S10), contrary to what is observed using a per-base mutation rate (Figure 3A). Indeed, when the genome-wide mutation rate is held constant, genome expansion no longer increases the mutational burden. Consequently, the protective benefit of increasing the number of gene copies is no longer offset by the increased risk of accumulating additional mutations. This demonstrates that assumptions about how mutation rates scale with genome size can fundamentally alter the evolutionary dynamics of genome expansion and robustness. Empirical estimates indicate that, for point mutations, the per-genome mutation rate increases with genome size (Lynch et al., 2023). Although similar data are not available for structural mutations, their rates are also expected to scale with genome size, which makes the assumption of a constant per-genome mutation rate un-realistic.

**Figure 4.**
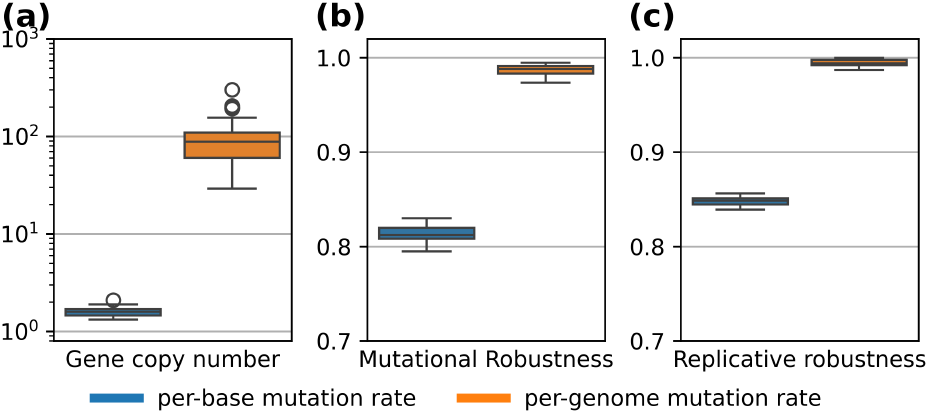
Comparison of (a) gene copy number, (b) mutational robustness and (c) replicative robustness at equilibrium for per-base mutation rates and per-genome mutation rates. The population size is fixed at 10, 000 to ensure efficient selection. For the per-genome mutation rates, rates are fixed at: duplication rate = 0.05, deletion rate = 0.15 and point mutation rate 0.5 per genome per replication. These rates were chosen to be similar to the per-genome mutation rates of the other experiments (per base duplication rate = 1 *×* 10^−7^, per base deletion rate = 3 *×* 10^−7^ and per base point mutation rate = 1 *×* 10^−6^; for a measured genome size of approximately 5 *×* 10^5^ bp at equilibrium.

## Sensitivity to parameter choices

To verify that our results are robust to specific assumptions, we have run versions of our model with different numbers of essential genes (50 here, also tested with 10, 5 and 1), different initial amounts of non-coding DNA between genes (0 in the presented experiments, also tested 200 and 2, 000), and different average sizes of rearrangements (here 1*/*10^*th*^ of the genome, we also tested 1*/*3^*rd*^ and 1*/*50^*th*^). We also ran a minimal model in which the genome is composed only of fully functional genes that can only be duplicated or deleted entirely (no point mutations, no non-functional DNA). Across all these simulations, our results remain qualitatively the same (see details and results in the Supplementary Materials S3: Figure S5, Figure S6 and Figure S7).

## Discussion

Selection for robustness is generally considered to reduce the deleterious impact of mutations. Our results show that selection can act on the number of mutations per replication, rather than only on their individual effects. Indeed, despite the absence of a direct cost for maintaining additional gene copies, and even though extra copies could reduce the risk of any single mutation being lethal, our model shows that gene redundancy is disfavoured across a wide range of circumstances. Instead, selection often favours maintaining the shortest possible genome, which minimises the number of mutational events per replication.

These results contrast with the classical “survival of the flattest” hypothesis (Wilke and Adami, 2003; Wilke et al., 2001), which predicts that selection favours genomes with greater robustness to deleterious mutations. The key to this discrepancy lies in the fact that mechanisms increasing mutational robustness often also increase genome size (*e*.*g*. via redundancy or expansion of non-coding regions), which in turn reduces replicative robustness. As we have shown, the net result can be that these mechanisms are in fact selected against. In terms of the traditional fitness landscape, this means that plateaus can be unstable (populations may drift off due to an increased risk of highly disruptive structural mutations), while peaks can be stable (populations are likely to stay there as the per-genome mutation rate is reduced).

The distinction we observe between mutational robustness and replicative robustness arises from two factors. First, mutations are more numerous in larger genomes (Figure 4). Second, chromosomal rearrangements are, on average, larger (and therefore more deleterious) as genome size increases. While there is evidence that rearrangements are indeed larger in organisms with larger genomes (Cui et al., 2012; da Silva et al., 2019; Wellenreuther and Bernatchez, 2018), the exact scaling relationship remains unknown. This relationship likely determines whether (and to what extent) mutational robustness and replicative robustness are positively or negatively correlated. In particular, the abundance and distribution of repeated elements may play a key role, as they can mediate chromosomal rearrangements (Schneider et al., 2000; Reams and Roth, 2015); their presence at distant loci could therefore reinforce the scaling between re-arrangement size and genome size. Overall, the precise relationship between genome size and the distribution of structural mutations is by no means elucidated. Broader empirical studies would help assess the extent to which the patterns observed in our study are likely to be relevant in real-world populations, and further theoretical work is needed to clarify the evolutionary consequences of different scaling regimes.

Our model has several limitations. We have not included an explicit cost of genome size, even though it might be realistic (*e*.*g*. due to replication time or nutrient constraints), as we wanted to avoid obfuscating the effects of selection due to robustness. Nevertheless, we expect that the main patterns of our results would still be there if such a cost were implemented - they would just be harder to interpret. Similarly, we have not considered other mechanisms that increase mutational robustness without affecting genome size, such as the distribution of connections and modularity of gene regulatory networks or the temporal stability of proteins (Goldstein, 2011; Hulst et al., 2025). In these cases, we hypothesise that replicative robustness would remain the primary target of robustness selection, but replicative and mutational robustness are likely to be much more intimately correlated.

Our results show that the traditional fitness landscape metaphor—where neighbouring genotypes differ by a single point mutation—can be misleading when considering robustness evolution. Indeed, selection acts on the outcome of replication events, which may include one or multiple mutations. An alternative conceptualisation in which neighbouring genomes are separated by full replication events may therefore be more appropriate, but risks becoming overwhelmingly complex. Visual metaphors may still be useful, such as oriented graphs where nodes are coloured by fitness and connected by the probability of going from one genome to another (the ‘replicative distance’). However, such a framework must accommodate complexities that cannot easily be simplified. In particular, transition probabilities between genomes are highly asymmetric. Because replication events can involve multiple, dependent mutations, the set of paths from genome A to genome B generally does not mirror those from B to A, making the two transition probabilities inherently unequal. This is especially true with structural mutations: a single deletion can remove a genomic segment in one step, whereas reconstructing the same sequence would require a series of highly specific duplications and point mutations, making the reverse transition astronomically less likely.

Our results also shed new light on the evolution of mutation rates. It illustrates that what selection ultimately sees is the per-genome mutation rate, which depends both on the per-base mutation rate and the genome size. While there is little doubt that the per-base mutation rate can be selected through changes in polymerase fidelity (Lynch et al., 2016), the per-genome mutation rate can also be reduced by compacting genomes (Luiselli et al., 2024). How these two main factors affecting replicative robustness are selected for, depending on the rates of structural and point mutations as well as other factors, remains an open avenue for further research.

An interesting direction for future work concerns the relationship between robustness and evolvability. Theory predicts that robustness in biological systems can lead to enhanced evolvability (Wagner, 2008b; Masel and Trotter, 2010; Hulst et al., 2025). For example, redundancy can promote evolvability by allowing duplicated elements to diverge, potentially leading to new functions. Hence, redundancy may increase evolvability while decreasing replicative robustness, creating a potential evolutionary trade-off between short-term transmission stability and long-term innovation potential. Extending our framework to include beneficial mutations, or environmental changes that alter the fitness associated with different variants, would allow this possibility to be explored explicitly. Such extensions could help explain why some clades evolve very large genomes (*e*.*g*., salamanders, plants) while others maintain very short genomes (*e*.*g*., marine cyanobacteria, fruit flies).

To conclude, our experiments show that while selection for robustness is indeed an important evolutionary force, it is replicative robustness, rather than robustness to single mutations, that is the primary target of selection. Because mutational robustness is a component of replicative robustness, the two may sometimes be selected in the same direction; however, there can also be a trade-off between them: a genome contraction can enhance replicative robustness at a cost to mutational robustness, as observed in our experiments. This also has implications for how we think about fitness landscapes, especially in the context of robustness evolution. In this context, the classical view of fitness landscape, in which neighbouring genotypes are separated by single mutations, can lead to misleading conclusions, particularly when considering the effect of fitness plateaus on the evolutionary dynamics. Indeed, when the neighbourhood is defined in terms of replication events rather than individual mutations, some plateaus can become peaks due to the high number of mutations each replication event entails: they are therefore much less neutral than they may appear at first sight.

## Data Availability Statement

The C++ code of the model is available on the KU Leuven GitLab: https://gitlab.kuleuven.be/u0183417/replormutrob. Results of the experiments, as well as notebooks to reproduce the figures of the paper, are available on Zenodo: https://zenodo.org/records/21128519.

## Acknowledgments

The authors would like to thank Paul Banse for fruitful discussions about the subject. The resources and services used in this work were provided by the VSC (Flemish Supercomputer Center), funded by the Research Foundation - Flanders (FWO) and the Flemish Government.

## Supplementary Information

### Supplementary Note S1: Temporal data corresponding to the Figures of the main text

We provide here the temporal data of the runs used for the Figures of the main text. This shows that all runs are at equilibrium at the time of the comparison (for which we take the average value between generations 14, 000 and 15, 000).

**A. Genome structure evolution for different point mutation rates**

**Figure S1.**
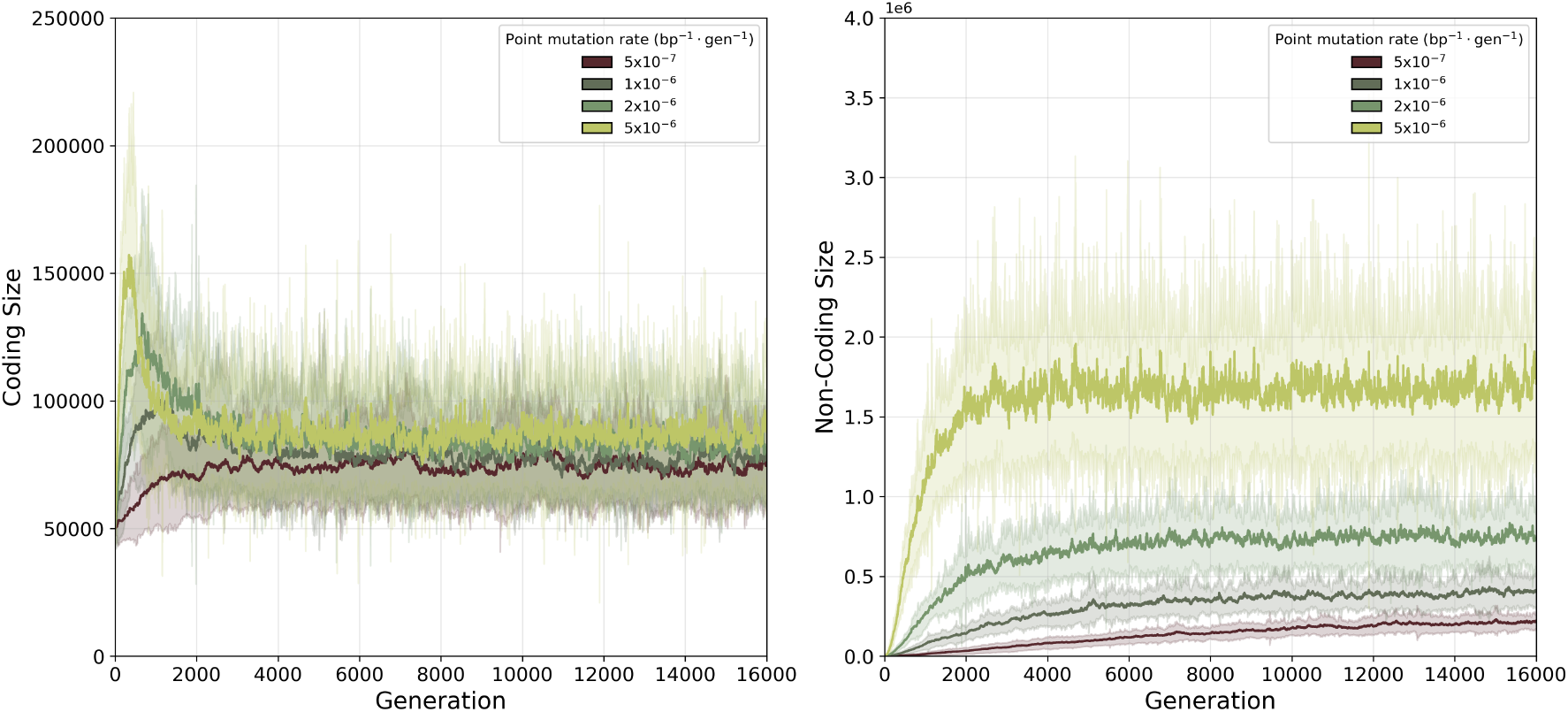
(A) Coding and (B) Non-coding size along the lineage of the ancestor of the final population, for the different point mutation rates, averaged over 50 replicates. Greyed areas show the standard deviation.

**B. Genome structure evolution for different deletion rates**

**Figure S2.**
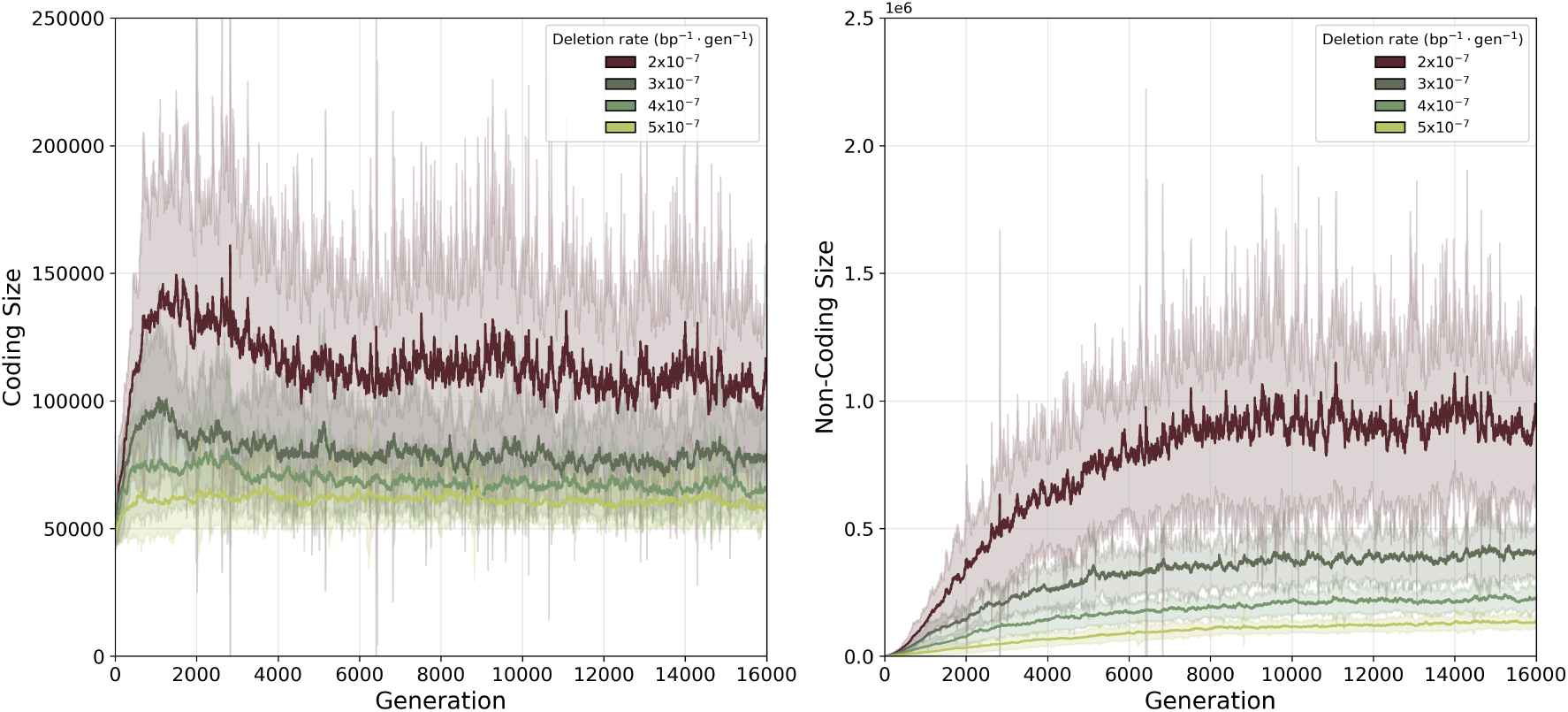
(A) Coding and (B) Non-coding size along the lineage of the ancestor of the final population, for the different deletion rates, averaged over 50 replicates. Greyed areas show the standard deviation.

**C. Genome structure evolution for different population sizes**

**Figure S3.**
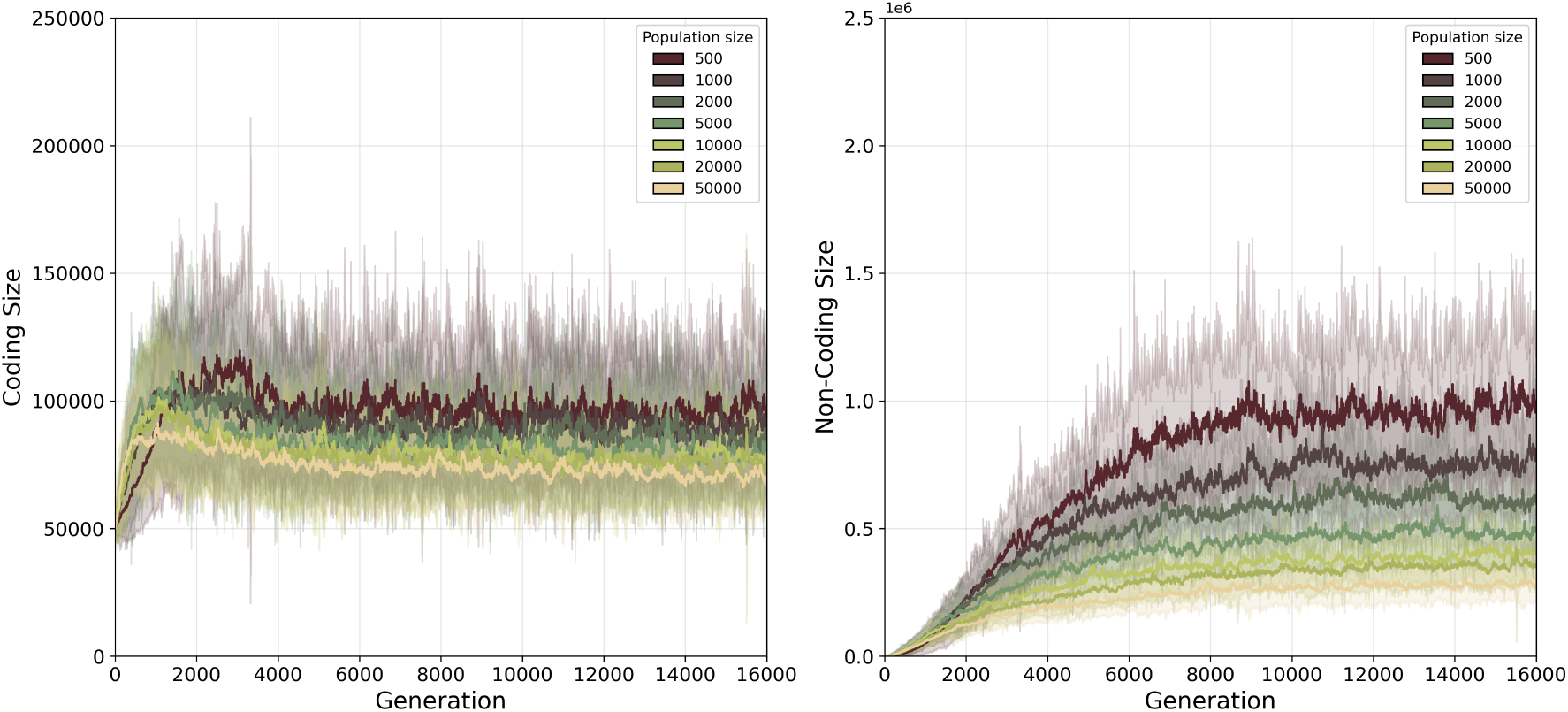
(A) Coding and (B) Non-coding size along the lineage of the ancestor of the final population, for the different population sizes, averaged over 50 replicates. Greyed areas show the standard deviation.

### Supplementary Note S2: Higher point mutation rates increase the non-coding genome size

**Figure S4.**
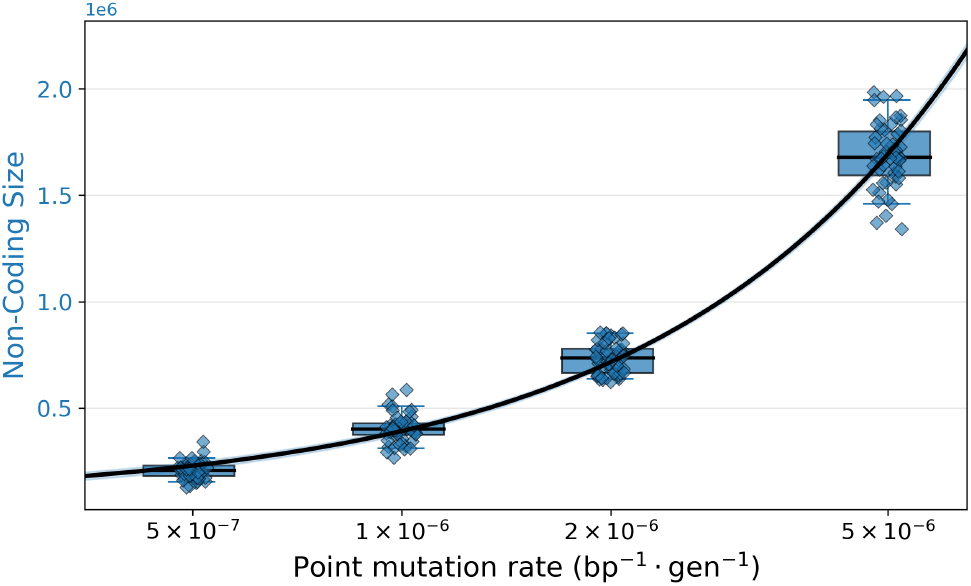
Amount of non-coding DNA (bp) for experiments with different point mutation rates. The other parameters (duplication rate = 1 *×* 10^−7^, deletion rate = 3 *×* 10^−7^, as well as population size *N* = 10, 000) are kept constant. A slight jitter has been added to the points to increase readability. Note that the scale of point mutation rate here is logarithmic to improve readability.

Due to the high rate of point mutations, genes are constantly pseudogenized, which increases the amount of non-coding DNA.

### Supplementary Note S3: Robustness of the model

#### A. Inversions

We tested the introduction of inversion mutations in our model, allowing genes to be reorganised along the genome. In particular, multiple copies of the same gene could be clustered together or dispersed within the genome. Inversions occur at a per-base rate, and their size follows the same distribution as duplications and deletions: a geometric law with an average of one-third of the genome size. This did not change the qualitative behavior of our model, as shown in the figures.

**Figure S5.**
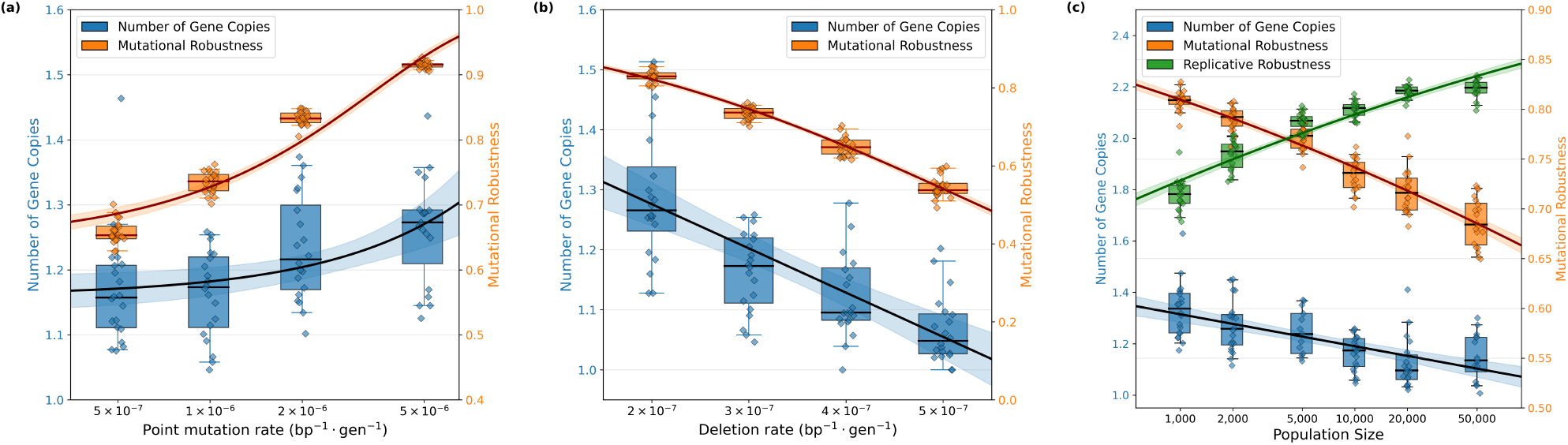
Average gene copy numbers (blue) and mutational robustness (orange) for different (a) point mutation rates, (b) deletion rates and (c) population sizes (red) with inversions added to the model. Data points are the average over generations 14, 000 to 15, 000 for the ancestor lineage of the final populations. Duplication rate is 1 *×* 10^−7^, inversion rate is 5 *×* 10^−7^. When not otherwise specified, point mutation rate is 1 *×* 10^−6^, deletion rate is 3 *×* 10^−7^ and population size is 10, 000.

#### B. Extreme simplification of the model

Model description We present here an extremely simplified version of our model: a genome is composed solely of genes, and mutations can only duplicate or delete full genes. As such, the mutation rate is per-gene, and the mutation length is measured in number of genes. Note that the average size of events is still proportional to the average size of the genome (here, the number of genes), and several genes can be duplicated/deleted at once. There is no non-functional genome in this model as genes cannot be inactivated, but solely entirely deleted. Genomes start with exactly one copy of each essential gene.

### Results

**Figure S6.**
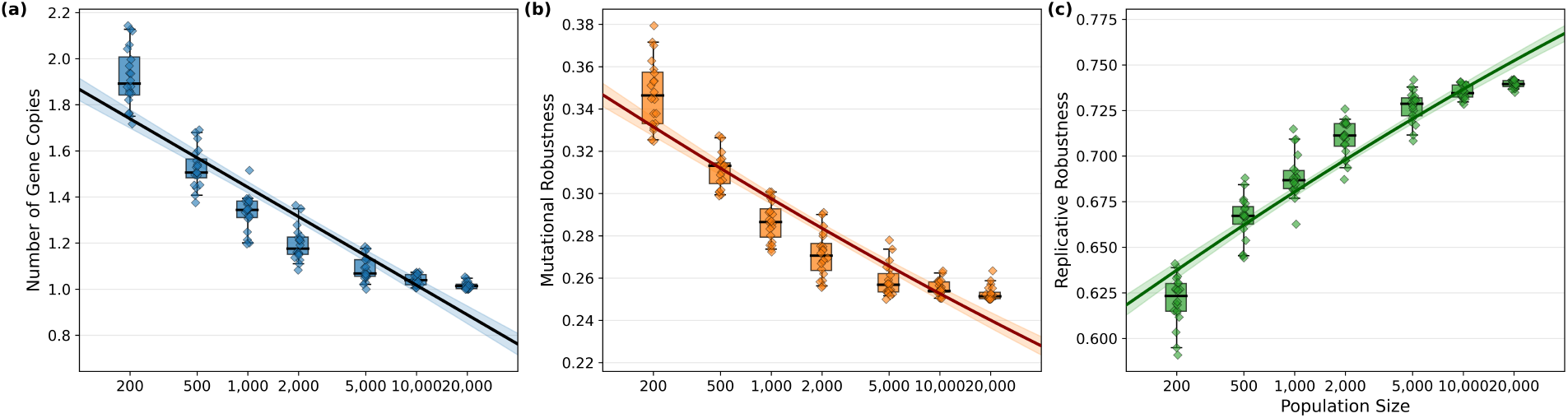
Average gene copy numbers (A), mutational robustness (B) and replicative robustness (C) for different population sizes, using the extremely simplified model. Data points are the average over generations 14, 000 to 15, 000 for the ancestor lineage of the final populations. Duplication rate is 10^−4^, and deletion rate is 3 *×* 10^−4^ per gene. There are no other types of mutations. 50 replicates per condition.

Even in this extremely simplified case, we still observe the same behaviour: a coding size reduction under larger population sizes. This shows that replicative robustness, rather than mutational robustness, is selected.

#### C. Different number of essential genes

Throughout the paper and the supplementary information, the number of essential genes was set to 50. We also tested lower numbers of essential genes (1, 5 and 10), while increasing the mutation rate to compensate for the reduced number of genes. For these tests, we used the simplified model (section B) to reduce computational cost, as preliminary analyses with fewer replicates showed that the simplified and the full model exhibit the same behaviour. Qualitatively, we observe the same behaviour regardless of the number of essential genes: The number of gene copies decreases as population sizes increases until it reaches a saturation point (corresponding to a minimum number of gene copies, see Fig. ??). However, this saturation point clearly depends on the number of essential genes. This observation can help quantitatively relate the model to actual biological data. In this example, we consider 50 genes and find that a population size of about 5, 000 individuals is sufficient to reach a state in which all essential genes are virtually present in a single copy. This suggests that larger (and more realistic) population sizes would be required to reach that same equilibrium when considering a larger (and more realistic) number of essential genes.

**Figure S7.**
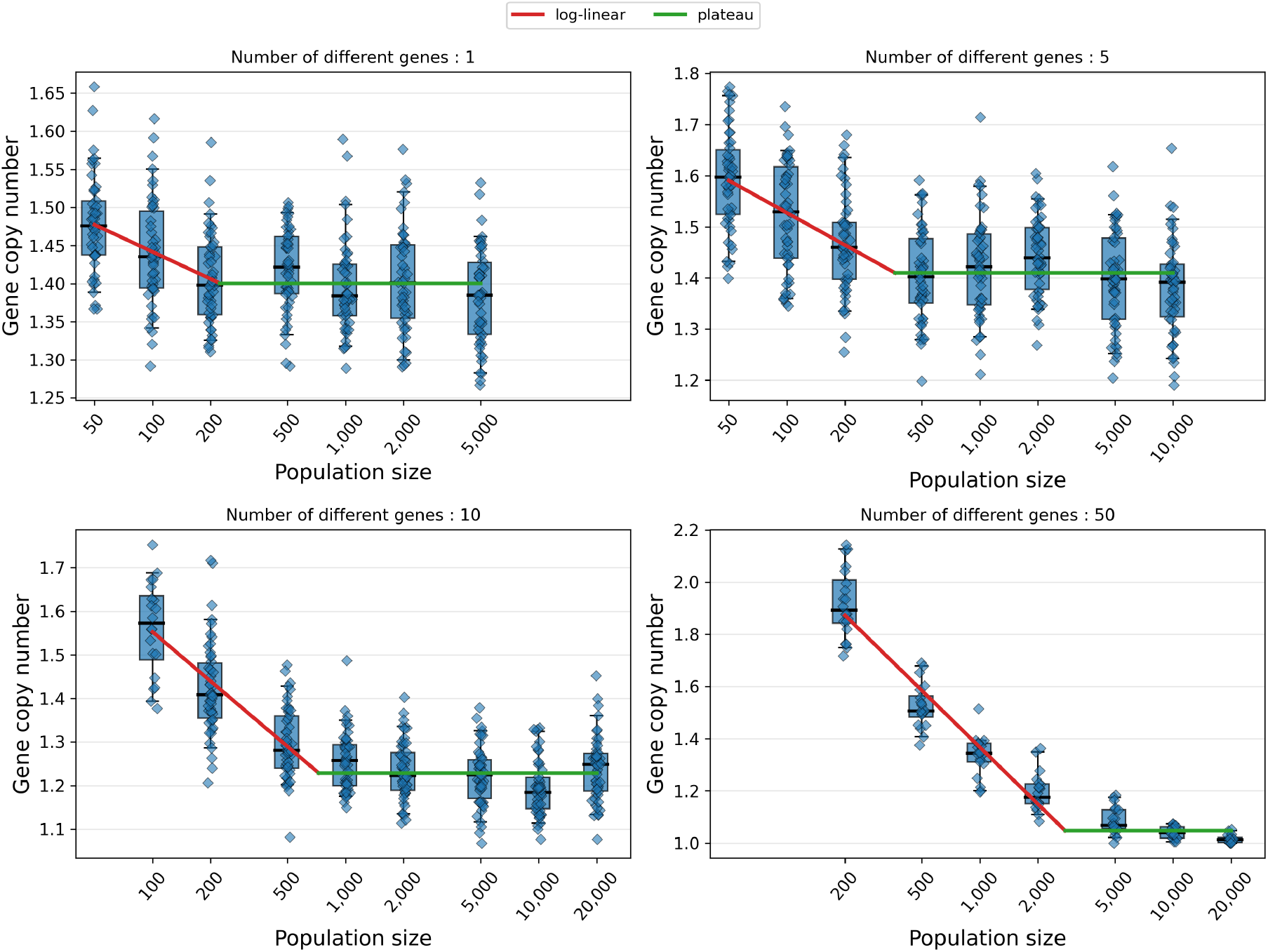
Average number of gene copies over generations 14, 000 to 15, 000 along the lineage of the ancestor of the final population, for different population sizes for (A) 1 gene, (B) 5 genes, (C) 10 genes and (D) 50 genes. For each panel, we fitted two linear curves to show the point of inflexion and the population size at which the optimum is reached. The mutation rates are: for 1 gene *µ*_dupl_ = 0.2bp−1 and *µ*_del_ = 0.6bp−1 ; for 5 genes *µ*_dupl_ = 0.02bp−1 and *µ*_del_ = 0.06bp−1 ; for 10 genes *µ*_dupl_ = 0.01bp−1 and *µ*_del_ = 0.03bp−1 and for 50 genes *µ*_dupl_ = 0.002bp−1 and *µ*_del_ = 0.006bp−1.

### Supplementary Note S4: Higher population size increases replicative robustness and decreases mutational robustness

**Figure S8.**
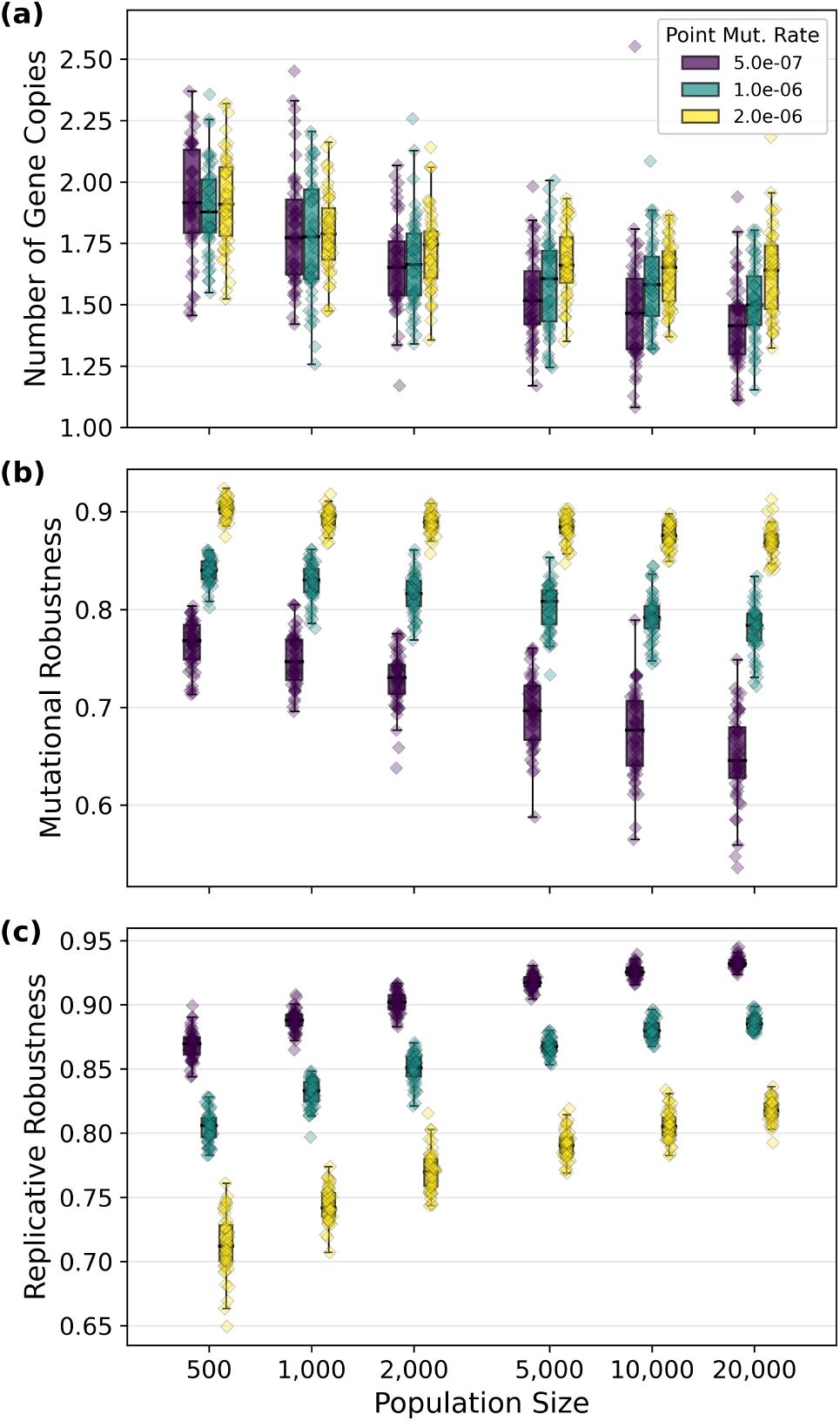
Average (a) number of gene copies (b) mutational robustness and (c) replicative robustness for various population sizes and point mutation rates. The other parameters are fixed at: duplication rate = 1 *×* 10^−7^, deletion rate = 3 *×* 10^−7^. 50 replicates per condition.

**Figure S9.**
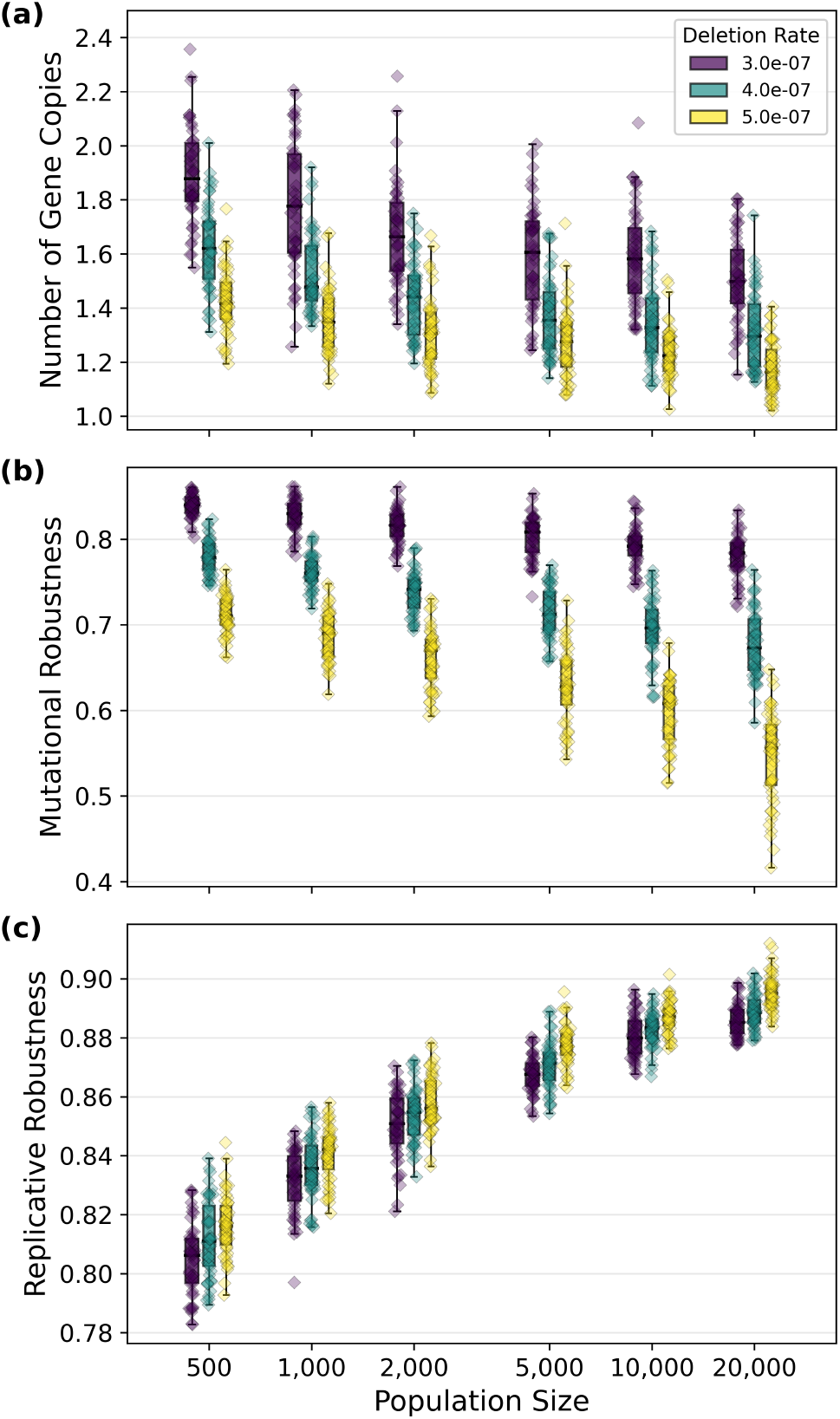
Average (a) number of gene copies (b) mutational robustness and (c) replicative robustness for various population sizes and deletion rates. The other parameters are fixed at: duplication rate = 1 *×* 10^−7^, point mutation rate = 1 *×* 10^−6^. 50 replicates per condition.

### Supplementary Note S5: Computational limit on genome size

During the experiments, some individuals can reach very high genome sizes by undergoing several large duplication events rapidly. Consequently, some individuals could have very high genome sizes and undergo a very large number of mutations, which caused the RAM to fill up and simulations to sometimes crash. However, these individuals generally died in the next generation due to the number of mutations, and were never part of the lineage of the final population if left to evolve. Therefore, a limit of 10^9^ bp per genome was implemented. Comparison of experiments run with and without this limit showed no difference in simulation output, but an important gain in time to run the simulations.

### Supplementary Note S6: Fixed per-genome mutation rate

**Figure S10.**
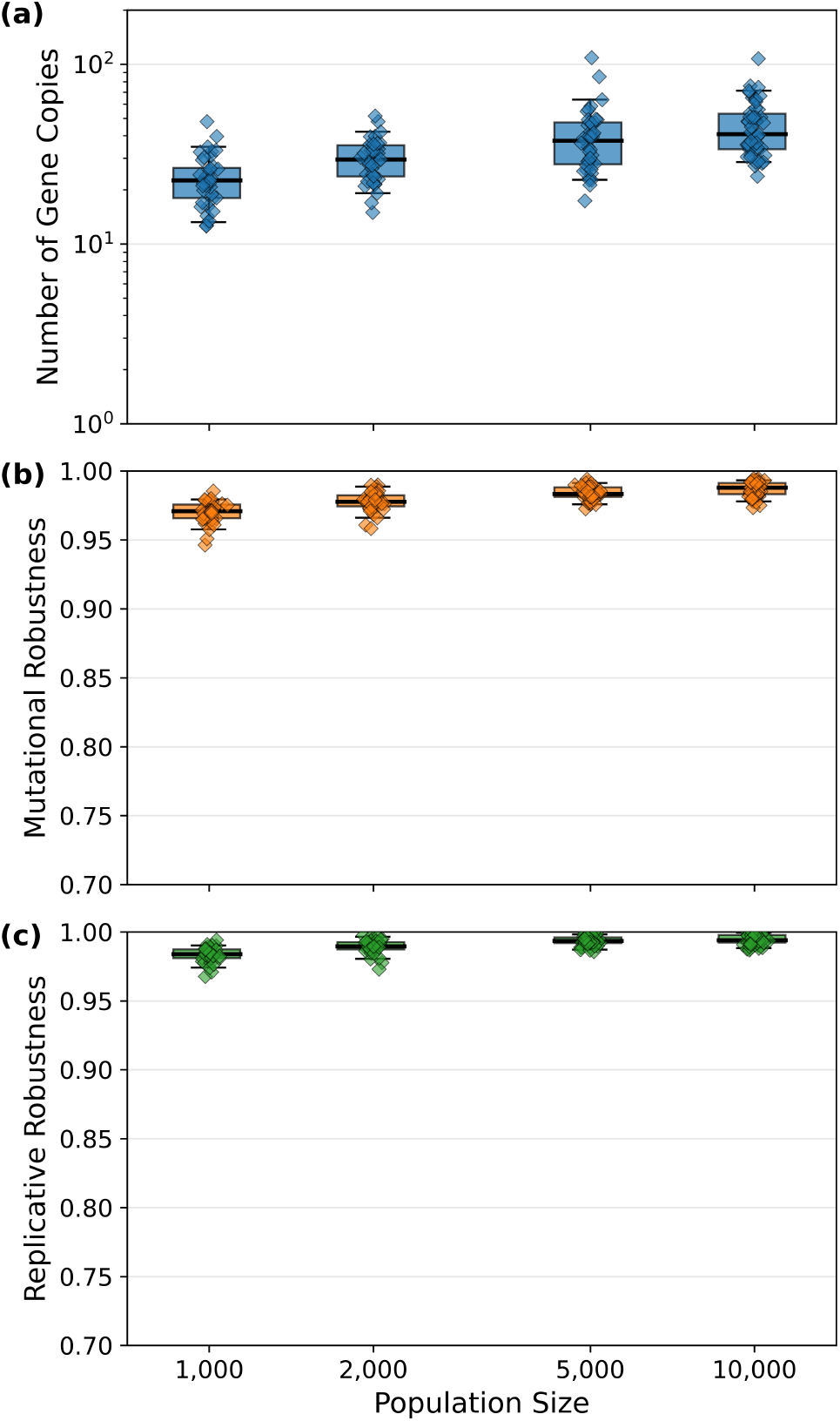
Average (a) number of gene copies (b) mutational robustness and (c) replicative robustness, for various population sizes. The other parameters are fixed at: duplication rate = 0.05, deletion rate = 0.15and point mutation rate 0.5 per genome per replication.

### Supplementary Note S7: Robustness as a function of genome size

We used the average genome structure evolved in our simulations of population size *N* = 10, 000, duplication rate of 1 × 10^−7^, deletion rate of 3 × 10^−7^ and point mutation rate of 1 × 10^−6^ (the evolved coding fraction is 16.6%) and generated random genomes that have the same structure but with the average number of gene copies varied from 1 to 2, with 100 independent genomes generated for each average copy number. On these randomly generated genomes, we perform 10, 000 point mutations, 10, 000 deletions, 10, 000 replications during which only point mutations can happen, 10, 000 replications during which only deletions can happen and 10, 000 replications during which both types of mutation can happen. We compute the average neutrality of these events (*i*.*e*. the proportion of mutations or replications that were neutral), which is the relevant measure of robustness.

**Figure S11.**
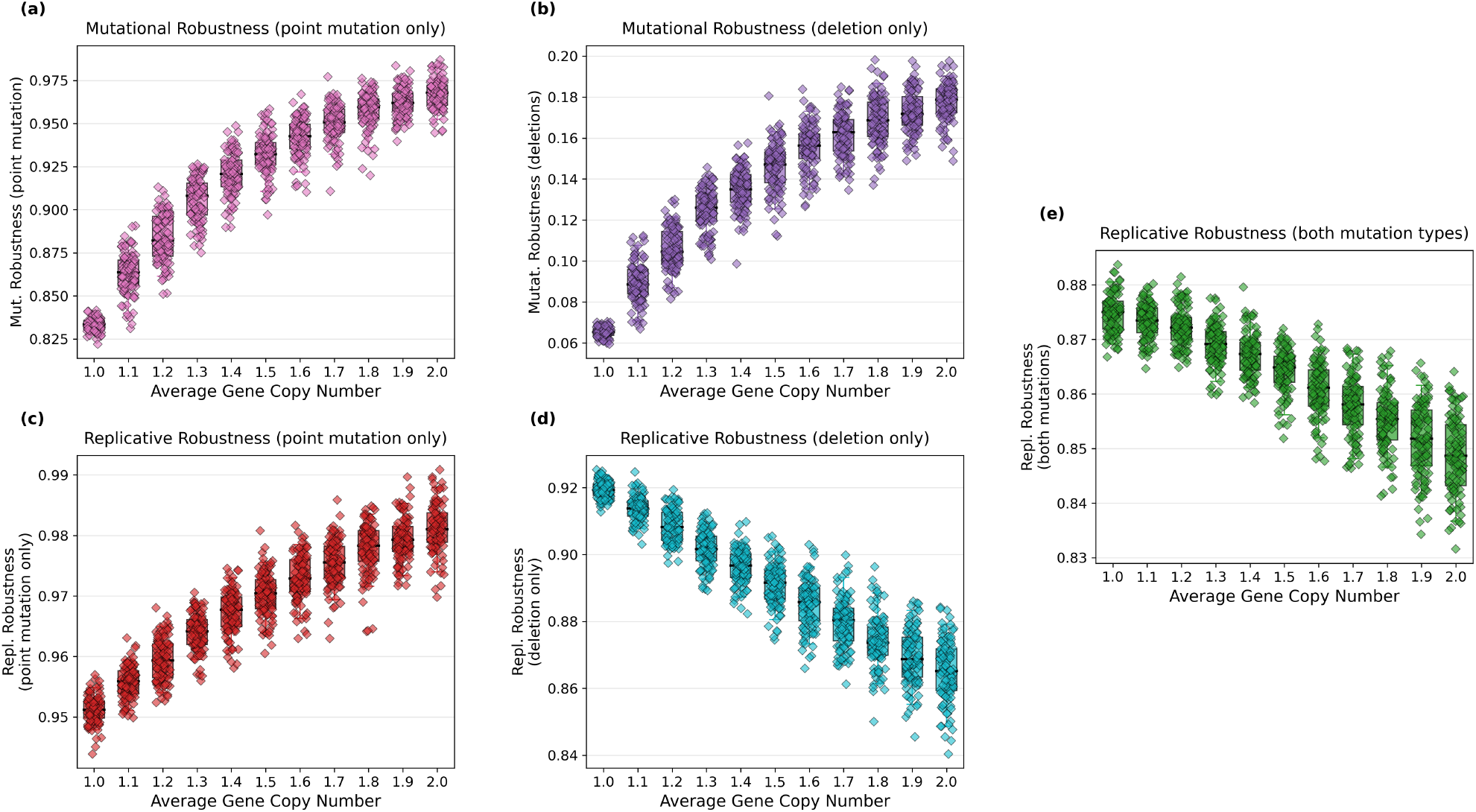
Measured robustness for different average gene copy numbers: (a) mutational robustness to a single point mutation, (b) mutational robustness to a single deletion, (c) replicative robustness when only point mutations can happen, (d) replicative robustness when only deletions can happen and (e) replicative robustness when both deletions and duplications can happen. This highlights the trade-off between mutational robustness and replicative robustness in the case of deletions: while the mutational robustness increases as the average number of gene copies increases, the replicative robustness decreases. As deletions are more dangerous than point mutations, they drive the total replicative robustness (panel e)

